## Supplementary material for "BDNF signaling in correlation-dependent structural plasticity in the developing visual system": Suppmenetal table 1

### **Supplemental Table 1**

| **Figure** | **F-statistic (interaction)** | **Friedman statistic** | **Kruskal-Wallis statistic** |
| --- | --- | --- | --- |
| 1F | **F(6,58)=2.78677; p=0.018884** | **Control: 8.667; p=0.0131**  TrkB-Fc: 3.308; p>0.1  p75-MO: 0.6667; p>0.1  TrkB-MO: 0.6087; p>0.1 | asynch: 1.406; p>0.1  **synch: 8.456; p=0.0375** |
| 1G | **F(6,58)=5.8315;**  **p<0.001** | Control: 4.979; p=0.083  TrkB-Fc: 5.407; p=0.0718  **p75-MO: 7.185; p=0.0212**  TrkB-MO: 4.571;p>0.1 | asynch: 3.690; p>0.1  **synch: 9.479; p=0.0236** |
| 2E | **F(6,56)=5.51030  p<0.001** | **Control:9.500; p=0.0087**  TrkB-Fc: 3.000; p>0.1  p75-MO: 0.8571; p>0.1  TrkB-MO:0.8571; p>0.1 | asynch: 4.945; p>0.1  **synch:7.994; p=0.0461** |
| 2F | F(6,58)=2.1958;  p=0.056209 |  |  |
| 3E | **F(6,102)=4.5683; p<0.001** | **Control: 25.07; p<0.0001**  **p75-MO: 17.59; p<0.001**  **TrkB-MO: 23.70; p<0.0001** | **Day 2: 7.969; p=0.0186**  Day 3: 5.461; p=0.0652  Day 4: 4.605; p=0.1 |
| 3F | **F(6,102)=9.2118;**  **p<0.001** | **Control: 41.00; p<0.0001**  **p75-MO: 14.76; p=0.002**  **TrkB-MO: 34.90; p<0.0001** | Day 2: 5.835; p=0.0541  **Day 3: 9.242; p=0.0098 Day 4: 9.984; p=0.0068** |
| 3G: bin1 (0-5 mm) | **F(6,102)=2.25385;**  **p=0.04394062** | **Control: 11.05; p=0.0114**  p75-MO: 1.848; p>0.1  TrkB-MO: 7.30; p=0.0629 | Day 1: 0.1906; p>0.1  Day 2: 5.197; p=0.0744  Day 3: 1.736; p>0.1  Day 4: 3.183; p>0.1 |
| 3G: bin2 (5-10 mm) | **F(6,102) = 2.6495;**  **p=0.019812** | **Control: 14.28; p=0.0025**  p75-MO: 6.152; p>0.05  TrkB-MO: 0.9000; p>0.1 | Day 1: 0.4411; p>0.1  **Day 2: 6.627; p=0.0364**  Day 3: 2.343; p>0.1  **Day 4: 6.571; p=0.0374** |
| 3G: bin3 (10-20 mm) | F(6,102)=0.96009; p>0.1 |  |  |
| 3G: bin4 (>20 mm) | F(6,102)=0.442131;  p>0.1 |  |  |
| 4D | **F(6,102)=4.0146;**  **P=0.0011918** | Control: 28.52; **p<0.0001**  p75-MO: 6.360; p=0.0954  TrkB-MO: 32.70; **p<0.0001** | Day1: 4.595; p>0.1  Day 2: 0.07752; p>0.1  Day 3: 2.683; p>p>0.1  Day 4: 3.072; p>0.1 |
| 4E |  |  | **8.086; p=0.0175** |
| S2A |  | **10.30; p=0.0357** |  |
| S2B |  | 6.889; p=0.1419 |  |
| S3A |  |  | 7.576; p=0.0556 |
| S3B |  |  | **8.631; p=0.0346** |
| S4A | **F(6,56)=7.1365;**  **p<0.001** | Control: 4.167; p>0.1  TrkB-Fc: 1.00; p>0.1 p75-MO: 3.174; p>0.1  TrkB-MO: 12.29; p<0.001 | asynch: 4.194; p>0.1  **synch: 9.5; p=0.0032** |
| S4B | **F(6,58)=6.9148;**  **p<0.001** | **Control: 8.167; p=0.0169**  TrkB-Fc: 3.714; p>0.1 p75-MO: 2.000; p>0.1  TrkB-MO: 3.714; p>0.1 | asynch: 7.541; p=0.0565  **synch: 12.73; p=0.0053** |
| S4C | **F(6,56)=5.79;**  **p<0.001** | **Control:11.17; p=0.0038**  TrkB-Fc: 3.00; p>0.1 p75-MO:1.143; p>0.1  TrkB-MO: 2.00; p>0.1 | asynch: 6.027; p>0.1  **synch: 10.66; p=0.0137** |
| S4D | **F(6,58)=3.2081; p=0.0086603** | Control:2.667; p>0.1  TrkB-Fc:3.429; p>0.1 p75-MO: 0.8571; p>0.1  TrkB-MO:0.2857; p>0.1 | asynch: 2.038; p>0.1  synch:6.888 ; p=0.0755 |
